# Diet-Induced Glial Insulin Resistance Impairs The Clearance Of Neuronal Debris

**DOI:** 10.1101/2023.03.09.531940

**Authors:** Mroj Alassaf, Akhila Rajan

## Abstract

Obesity significantly increases the risk of developing neurodegenerative disorders, yet the precise mechanisms underlying this connection remain unclear. Defects in glial phagocytic function are a key feature of neurodegenerative disorders, as delayed clearance of neuronal debris can result in inflammation, neuronal death, and poor nervous system recovery. Mounting evidence indicates that glial function can affect feeding behavior, weight, and systemic metabolism, suggesting that diet may play a role in regulating glial function. While it is appreciated that glial cells are insulin sensitive, whether obesogenic diets can induce glial insulin resistance and thereby impair glial phagocytic function remains unknown. Here, using a *Drosophila* model, we show that a chronic obesogenic diet induces glial insulin resistance and impairs the clearance of neuronal debris. Specifically, obesogenic diet exposure downregulates the basal and injury-induced expression of the glia-associated phagocytic receptor, Draper. Constitutive activation of systemic insulin release from *Drosophila* Insulin-producing cells (IPCs) mimics the effect of diet-induced obesity on glial draper expression. In contrast, genetically attenuating systemic insulin release from the IPCs rescues diet-induced glial insulin resistance and draper expression. Significantly, we show that genetically stimulating Phosphoinositide 3-kinase (PI3K), a downstream effector of Insulin receptor signaling, rescues HSD-induced glial defects. Hence, we establish that obesogenic diets impair glial phagocytic function and delays the clearance of neuronal debris.

## Introduction

Obesity significantly increases the risk for developing neurodegenerative disorders [1–3], yet the precise mechanisms underlying this connection remain unclear. Overconsumption of high sugar foods is the leading cause of obesity and its comorbidities including type 2 diabetes [4]. These effects are largely mediated by the breakdown of the insulin signaling pathway. Upon chronic exposure to a high-sugar diet (HSD), circulating insulin levels rise, resulting in insulin resistance, a state characterized by reduced cellular responsiveness to insulin [5]. Although HSD-induced insulin resistance has been well documented to occur in peripheral organs such as fat tissue and the liver, less is known about whether HSD-induced insulin resistance also occurs in the brain. Despite the widespread expression of insulin receptors in the brain, the brain has been historically regarded as insulin insensitive. This is largely because insulin is dispensable for glucose uptake in the brain [6] and its entry is limited by the blood-brain barrier [7, 8]. However, growing evidence suggest that insulin exerts unique regulatory actions on the brain to control cognition, feeding, and systemic metabolism[9, 10]

A key feature of neurodegenerative disorders is the diminished clearance of neuronal debris and neuron-secreted toxic proteins [11]. This can lead to inflammation, secondary neuronal death, and impaired axonal regeneration. Microglia are the brain’s resident macrophages, and they play a key role in learning and memory. When activated, they can swiftly mobilize to the site of disease or neuronal injury and initiate phagocytosis [12]. However, chronic activation of microglia can lead to the progressive decline of their phagocytic capacity as seen in the aging brain—the most at risk for neurodegenerative disorders [13]. Interestingly, obese humans and animals also display chronic activation of microglia, which has been shown to contribute to neuroinflammation [14]. However, little is known about the effects of obesity on glial phagocytic function. Uncovering whether and how diet-induced obesity disrupts glial phagocytosis may shed some light on the link between obesity and neurodegenerative disorders.

It was only recently discovered that microglia express insulin receptors indicating that insulin can have a direct regulatory effect on microglial function [15, 16]. While these studies suggest a tight link between glial cell function and insulin, precisely how physiological factors like diet-induced alterations in insulin levels modulate glial function remains unclear.

Glial phagocytosis begins with the recognition of cellular debris via cell-surface receptors. Ablation of these receptors results in impaired clearance of cellular debris, while their overexpression leads to excessive neuronal pruning. Just like in mammals, *Drosophila* microglia-like cells, ensheathing glia [17], express phagocytic receptors, most prominently the mammalian Multiple EGF Like Domains 10 (MEGF10) homolog, Draper [18]. Several studies have demonstrated that baseline levels of Draper in the uninjured brain determine the phagocytic capacity of ensheathing glia [19, 20]. Upon injury or disease, ensheathing glia upregulate Draper[19, 21–23]. However, low baseline levels may prevent Draper reaching a critical threshold for target detection leading to impaired clearance. Interestingly, Draper’s baseline levels were found to be regulated by Phosphoinositide 3-kinase (PI3K), a downstream effector of Insulin receptor (IR) signaling, while injury-induced Draper upregulation is regulated by another insulin signaling downstream target, the transcription factor Stat92E [19]. While local glial insulin receptor signaling has been shown to be a key regulator of Draper expression [24], it remains unknown whether obesogenic diets disrupt insulin signaling in glia and whether that disrupted signaling affects Draper expression and glial function. Hence, we set out to address whether prolonged obesogenic diets in *Drosophila* disrupt glial phagocytic function. We have previously shown that prolonged high sugar diet (HSD) treatment causes peripheral insulin resistance in adult flies [25]. Here, we show that chronic HSD exposure leads to insulin resistance in ensheating glia, which results in their impaired ability to clear axotomized olfactory neurons. Genetically inducing insulin release recapitulates HSD-induced Draper downregulation, while attenuating Insulin release rescues HSD-induced Draper downregulation. Importantly, we show that genetically stimulating a downstream effector of Insulin Receptor signaling in ensheathing glia rescues HSD-induced insulin resistance and the downregulation of Draper. Together, this study provides the first *in vivo* evidence of diet-induced regulation of glial phagocytic function.

## Results

### HSD affects the brain’s metabolism and causes lipid droplet accumulation

Our lab established that adult *Drosophila* subjected to a prolonged (> 2 weeks) high sugar diet (HSD-see methods) display hallmarks of peripheral insulin resistance including disrupted hunger response [25]. Using the obesogenic diet paradigm we established, we sought to characterize the effects of prolonged HSD treatment on the brain. All cells rely on two main sources of energy: glycolysis (a series of cytosolic biochemical reactions to generate ATP) and mitochondrial oxidative phosphorylation (OxPhos); OxPhos and glycolysis exist in a delicate balance [26, 27] (Figure 1A). To get a broad understanding of the effects of HSD on the brain’s glycolytic state, we sought to examine the expression levels of the glycolytic enzyme, lactate dehydrogenase (Ldh). Given that Ldh is responsible for the final step of glycolysis, it has been used as a reliable readout of glycolysis [28], especially using an LDH reporter in flies [29, 30]. We subjected a transgenic fly line that expresses a fluorescent Ldh reporter to either ND or HSD for 3 weeks. For this study, we chose to focus on the antennal lobe region given its accessibility and well-defined histology. The HSD-fed flies had significantly lower levels of Ldh in the antennal lobe region compared to the ND-fed flies (Figure 1B-C) suggesting attenuated glycolysis.

**Figure 1.**
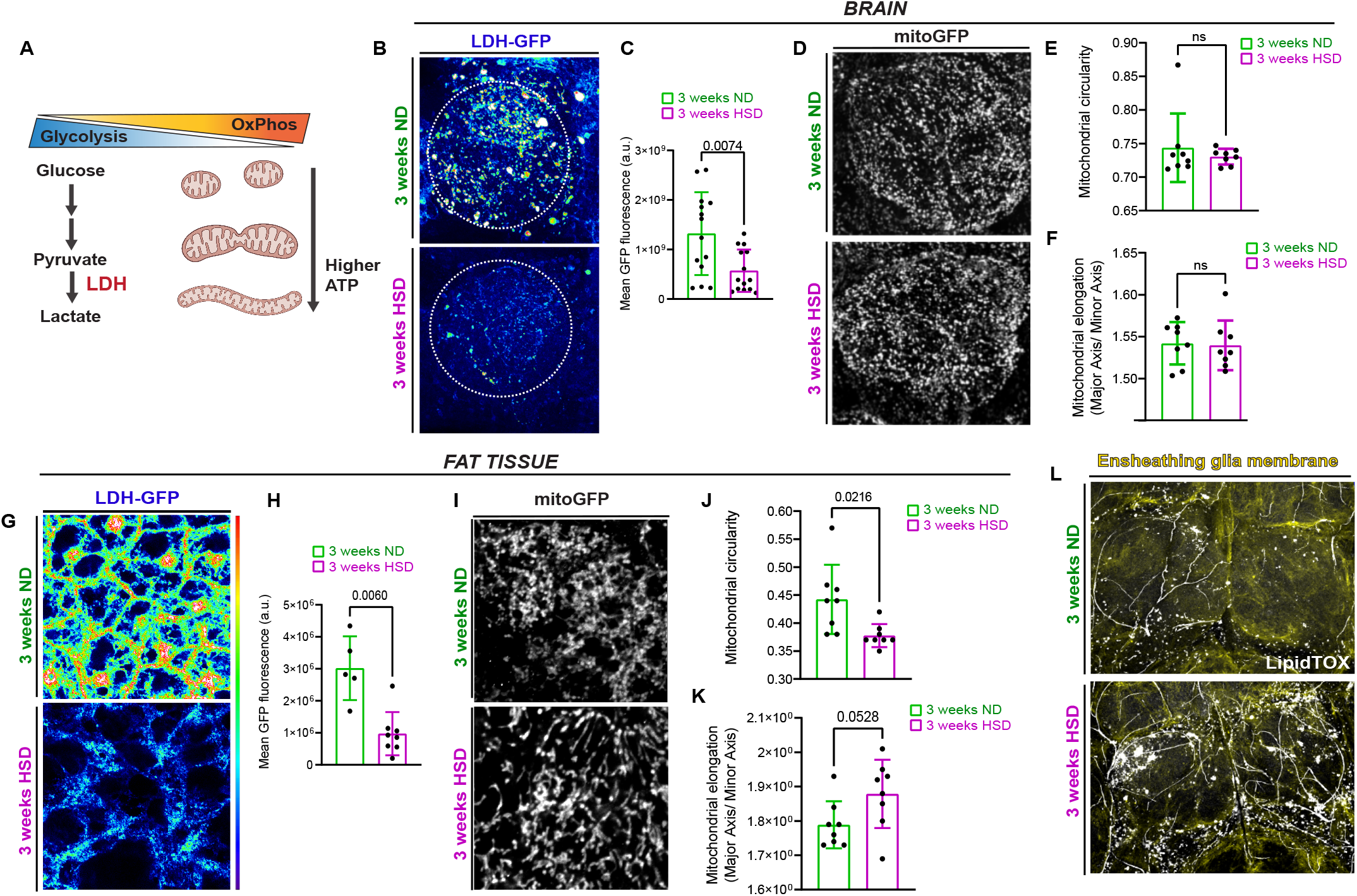
HSD affects the brain’s metabolism and causes lipid droplet accumulation. **A)** Schematic showing the cellular energy balance between glycolysis and mitochondrial oxidative phosphorylation (OxPhos). Elongated mitochondria are associated with increased OxPhos. **B)** Confocal images of LDH-GFP in the antennal lobe region (dotted circle) of ND and HSD-fed flies. **C)** Mean fluorescent intensity of GFP measured within a region of interest (dotted circle) that covers the antennal lobe in ND and HSD-fed flies. **D)** Confocal images of GFP-tagged mitochondria in the antennal lobe region of ND and HSD-fed flies. **E-F)** Mean mitochondrial (E) circularity and (F) elongation measured from Z-stack max intensity projections of the antennal lobe region of ND and HSD-fed flies. **G)** Confocal images of LDH-GFP in the fat tissue of ND and HSD-fed flies. **H)** Mean fluorescent intensity of GFP measured in the fat tissue of ND and HSD-fed flies. **I)** Confocal images of GFP-tagged mitochondria in the fat tissue of ND and HSD-fed flies. **J-K**) Mean mitochondrial (J) circularity and (K) elongation measured from Z-stack max intensity projections of the fat tissue of ND and HSD-fed flies. Student’s T test. N= each circle represents an individual fly. **L)** Confocal images of Lipitox-stained (white) lipid droplets in the brains of ND and HSD-fed flies that express membrane-tagged GFP in their ensheathing glia (yellow).

Typically, when glycolysis is stunted, the mitochondria undergo rapid expansion to compensate for the fall in energy levels (Figure 1A) [26, 27, 31]. Given that glycolysis occurs primarily in glia [32–35], we sought to investigate the effects of HSD on glial mitochondria. Specifically, we focused on ensheathing glia, which are functionally similar to microglia and reside within the antennal lobe [22]. As it is technically challenging to measure oxygen consumption rate[36] – a more direct readout for mitochondrial OxPhos-in intact glia without also obtaining readouts from the neurons, we took an imaging-based approach. Mitochondrial morphology can be used as a readout of their activity. The OxPhos capacity of elongated mitochondria is higher than that of circular mitochondria and is used as an accepted measure to indicate OxPhos capacity [31, 37–39]. Hence, we expressed a mitochondrion targeted GFP (mito-GFP) in ensheathing glia and analyzed mitochondrial morphology. We expected that since LDH expression was reduced we would observe elongated mitochondria. Surprisingly, although 3 week of HSD exposure reduced LDH reporter expression in the brain suggesting reduced glycolysis (Figure 1B-C), it did not alter mitochondrial morphology in ensheathing glia as assessed by circularity and elongation (Figure 1D-F). In contrast, in adult *Drosophila* abdominal adipocytes, we observe a clear shift in mitochondrial morphology to appear more elongated/less circular (Figure I-K) in response to the reduced LDH levels (Figure 1G-H) of HSD-fed flies. We interpret this to mean that the brain and adipocytes respond differently to prolonged HSD at 3-weeks. Based on this, we postulate that the adipocytes maintain their ability, even on HSD, to shift towards OxPhos in response to reduced glycolysis, but the glial cells are unable to do so (See Discussion).

Human and animal studies have shown that microglia accumulate lipid droplets in aging and neurodegenerative disorders, leading to impaired function [40, 41]. We have previously shown that prolonged HSD treatment increases the number and size of lipid droplets in adipocytes [25]. Thus, we asked whether HSD has a similar effect on glial lipid storage. To answer this, we visualized the lipid droplets in the brains of ND and HSD-fed flies using LipidTOX, a neutral lipid stain, and drove membrane-tagged GFP expression specifically in ensheathing glia. We found that prolonged HSD treatment markedly increased the number and size of lipid droplets in ensheathing glia surrounding the antennal lobes (Figure 1I) indicating possible glial dysfunction.

### HSD causes insulin resistance in glia

Lipid droplet accumulation and reduced glycolysis are tightly associated with impaired insulin signaling [25, 42–45]. Though obesogenic diets have been established by us and others to cause peripheral insulin resistance [25, 46, 47], whether obesogenic diets lead to central brain insulin resistance remains unclear. In a previous study, our lab demonstrated that chronic HSD treatment of 2 weeks or more causes insulin resistance in the adult *Drosophila* fat tissue [25]. To determine whether HSD treatment causes insulin resistance in the brain, we first compared the levels of the *Drosophila* Insulin-like peptide 5 (Dilp5) retained in the insulin producing cells (IPCs) of ND and HSD-fed flies. Dilp5 is primarily produced by the IPCs and its secretion is dictated by nutrient abundance [48, 49]. Dilp5’s retention in the IPCs is often used as a readout of its secretion [25, 50–55]. As expected, the HSD-fed flies showed reduced Dilp5 accumulation in their IPCs suggesting increased Dilp5 secretion (Figure 2A-B).

**Figure 2.**
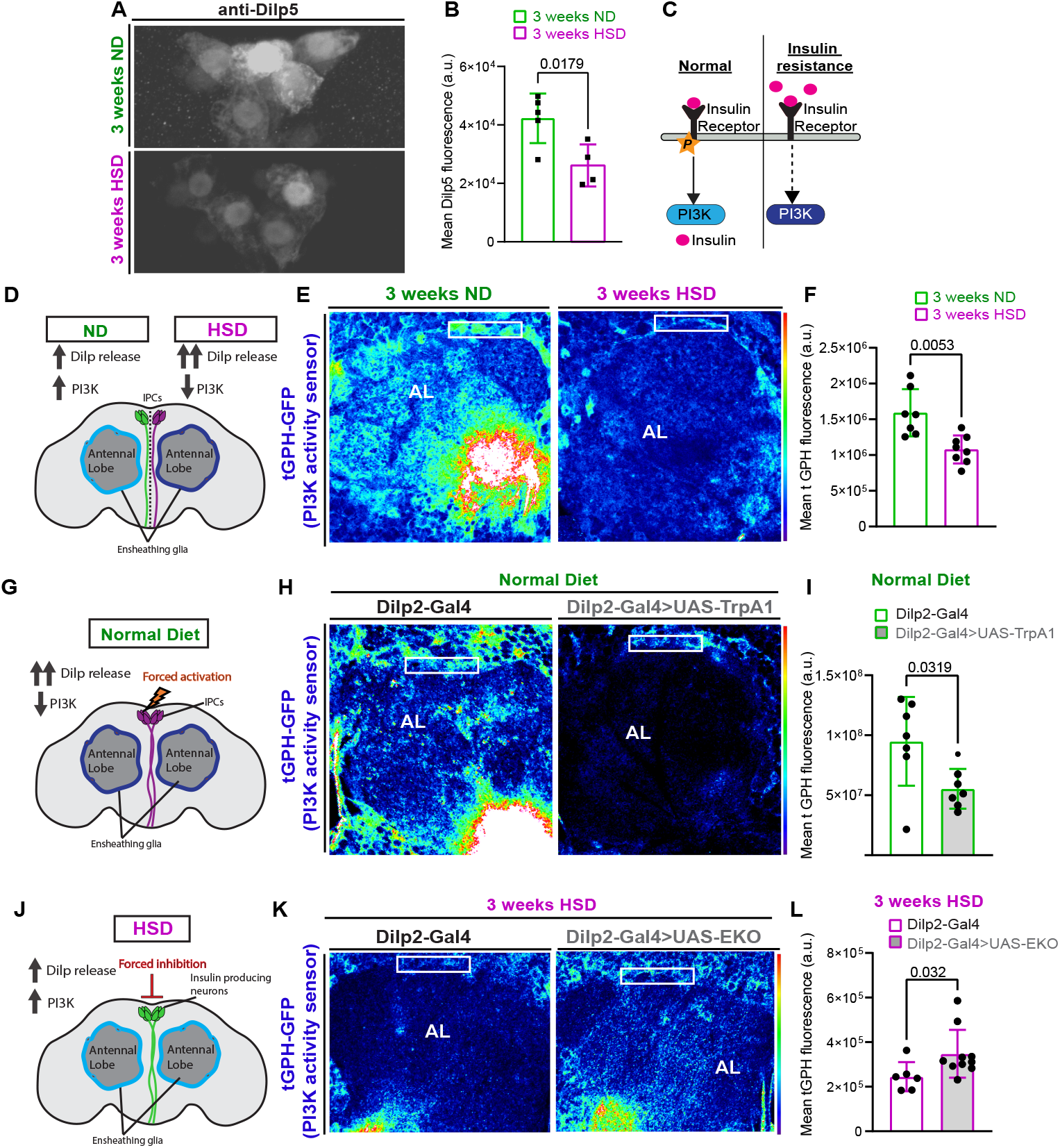
HSD causes insulin resistance in glia. **A)** Representative confocal images of Dilp5 accumulation in the insulin-producing cells (IPCs) of ND and HSD-fed flies. **B)** Mean fluorescent intensity of anti-Dilp5 in the IPCs of ND and HSD-fed flies. **C)** Schematic of the insulin resistance model. Increased circulating insulin desensitizes the insulin receptors leading to reduced PI3K activity. **D)** Schematic for experimental design and summarized results of (E-F). **E)** Confocal images of the antennal lobe region of flies expressing the PI3K (tGPH) activity sensor that were either fed a ND or a HSD. **F)** Mean fluorescent intensity of GFP measured within a region of interest (white box) that coincides with the location of ensheathing glia in ND and HSD-fed flies. **G)** Schematic for experimental design and summarized results of (H-I). **H)** Confocal images of tGPH levels in the antennal lobe region of control flies and flies expressing TrpA1 in the IPCs that were fed a ND. TrpA1 expression force activates the IPCs to release Dilps. **I)** Mean fluorescent intensity of GFP measured within a region of interest (white box) that coincides with the location of ensheathing glia in control and flies with Dilp2-driven TrpA1 expression. **J)** Schematic for experimental design and summarized results of (K-L). **K)** Confocal images of tGPH levels in the antennal lobe region of control flies and flies expressing EKO in the IPCs that were fed a HSD. EKO expression attenuates the release of Dilps from the IPCs. **L)** Mean fluorescent intensity of GFP measured within a region of interest (white box) that coincides with the location of ensheathing glia in control and flies with Dilp2-driven EKO expression. Student’s T test. N= each circle represents an individual fly.

To investigate whether the increase in insulin signaling led to glial insulin resistance, we used a transgenic line that expresses a fluorescent reporter (tGPH) for Phosphoinositide 3-kinase (PI3K) [56, 57], a downstream effector of Insulin receptor (IR) signaling. Under normal conditions, the IR autophosphorylates upon interacting with insulin, which leads to activation of the PI3K pathway. However, excessive levels of circulating insulin can attenuate IR sensitivity leading to reduced PI3K activation [58, 59] (Figure 2C). To this end, we subjected the tGPH flies to 3 weeks of either ND or HSD and measured tGPH fluorescence in the area surrounding the antennal lobe where ensheathing glia reside (Figure 2D-E). We found that HSD caused a significant downregulation of PI3K activity indicating insulin resistance (Figure 2D-F). Notably, we did not observe any gross morphological defects in response to the prolonged HSD treatment (Figure S1). To determine whether the HSD-induced attenuation of PI3K signaling is specifically due to excessive systemic insulin secretion, we genetically induced insulin secretion from the IPCs by expressing the neuronal activator TrpA1 under the control of an IPC-specific Gal4 driver [51, 55]. Remarkably, we found that forced activation of the IPCs for 1 week mimics the effects of a 3-week HSD exposure on PI3K activity in ensheathing glia (Figure 2G-I). In contrast, attenuating the release of insulin in the HSD-fed flies by expressing a genetically modified potassium channel (EKO) that inhibits neuronal activation [60, 61] under the control of an IPC-specific driver increases glial insulin signaling (Figure 2J-L). Together, these findings demonstrate that obesogenic diets directly cause glial insulin resistance through excess systemic insulin.

### HSD downregulates basal Draper levels

As the brain’s resident macrophages, microglia are crucial to the survival and function of the nervous system through their phagocytic activity [62]. Just like the mammalian microglia, the phagocytic activity of ensheathing glia is governed by the engulfment receptor, Draper (The *Drosophila* ortholog to the mammalian MEGF10) [22, 23]. Given that PI3K signaling has been shown to regulate basal Draper levels [19], we reasoned that HSD treatment and the ensuing downregulation of PI3K signaling (Figure 2D-F) would result in reduced basal Draper levels. To test this, we measured Draper immunofluorescence within a subset of ensheathing glia in ND and HSD-flies. We found that 3 weeks of HSD treatment caused a substantial reduction in basal Draper levels (Figure 3A-C).

**Figure 3.**
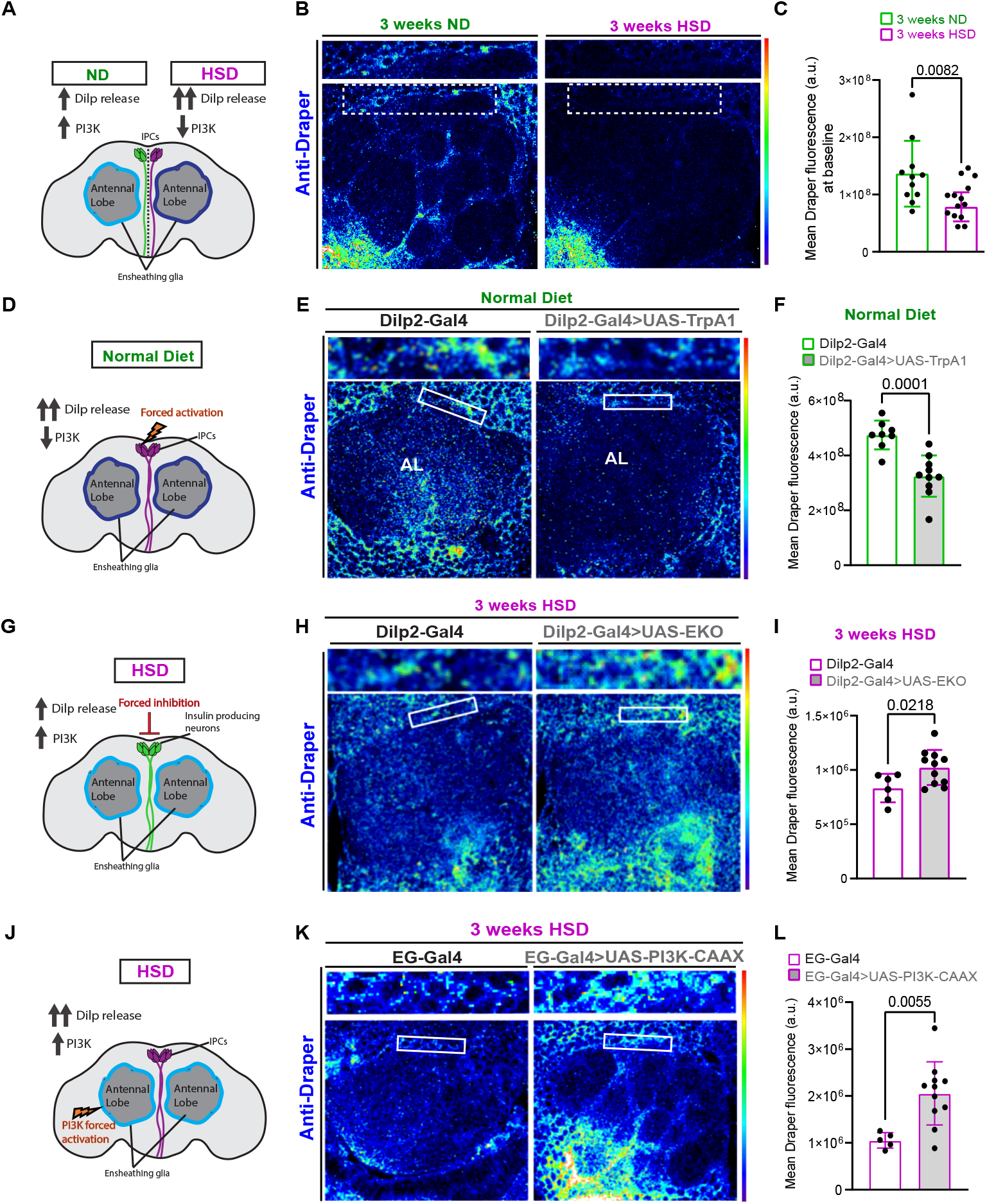
HSD downregulates basal Draper levels. **A)** Schematic for experimental design and summarized results of (B-C). **B)** Confocal images of the antennal lobe region of flies fed a ND or a HSD that were immunostained with anti-Draper. **C)** Mean fluorescent intensity of Draper measured within a region of interest (white box) that coincides with the location of ensheathing glia in ND and HSD-fed flies. **D)** Schematic for experimental design and summarized results of (E-F). **E)** Confocal images of Draper levels in the antennal lobe region of control flies and flies expressing TrpA1 in the IPCs that were fed a ND. TrpA1 expression force activates the IPCs to release Dilps. **F)** Mean fluorescent intensity of Draper measured within a region of interest (white box) that coincides with the location of ensheathing glia in control and flies with Dilp2-driven TrpA1 expression. **G)** Schematic for experimental design and summarized results of (H-I). **H)** Confocal images of Draper levels in the antennal lobe region of control flies and flies expressing EKO in the IPCs that were fed a HSD. EKO expression attenuates the release of Dilps from the IPCs. **I)** Mean fluorescent intensity of Draper measured within a region of interest (white box) that coincides with the location of ensheathing glia in control and flies with Dilp2-driven EKO expression. **J)** Summarized results of (K-L). **K)** Confocal images of Draper levels in the antennal lobe region of HSD-fed control flies and flies expressing PI3K-CAAX in ensheathing glia. **L)** Mean fluorescent intensity of Draper measured within a region of interest (white box) that coincides with the location of ensheathing glia. Student’s T test. N= each circle represents an individual fly.

It is possible that HSD treatment results in non-insulin dependent downregulation of Draper signaling. Therefore, we reasoned that if glial insulin resistance (Figure 2) is responsible for Draper downregulation in HSD-fed flies, then forced systemic insulin release from the IPCs would also result in Draper downregulation. Indeed, we find that expressing the neuronal activator TrpA1 under the control of an IPC-specific Gal4 driver leads to reduced Draper levels under ND conditions (Figure 3D-F). In contrast, attenuating systemic insulin release from the IPCs by expressing a genetically modified potassium channel (EKO) that inhibits neuronal activation paradoxically increases Draper expression in the HSD-fed flies (Figure 3G-I).

Next, we reasoned that if glial insulin resistance underlies the downregulation of Draper in HSD-fed flies, then stimulating PI3K signaling, a downstream arm of insulin signaling, will upregulate Draper expression. To address this, we expressed a constitutively active form of PI3K that is fused to a farnesylation signal (CAAX) [63] under the control of an ensheathing glia promoter. We found that stimulating ensheathing glia PI3K signaling increases Draper expression under HSD compared to the HSD-fed controls (Figure 3J-L). Together, this suggests that HSD downregulates Draper expression by inducing glial insulin resistance.

### HSD delays the clearance of degenerating axons by inhibiting injury-induced Draper and STAT upregulation

Normally, neuronal injury triggers the upregulation of Draper in ensheathing glia that peaks one day after injury and persists until neuronal debris has been cleared [23]. Therefore, we asked whether HSD treatment prevents the upregulation of Draper after neuronal injury. To answer this, we took advantage of the accessibility of the olfactory neurons. We performed unilateral ablation of the third antennal segment, which houses the cell bodies of olfactory neurons. This results in the Wallerian degeneration of olfactory neurons’ axons that project to the antennal lobe, which induces ensheathing glia to phagocytose axonal debris [22, 23, 64]. Then, we immunostained for Draper one day post antennal ablation (Figure 4A). As expected, Draper levels increased significantly in the ND-fed flies, whereas the HSD-fed flies showed no upregulation in Draper (Figure 4B-C).

**Figure 4.**
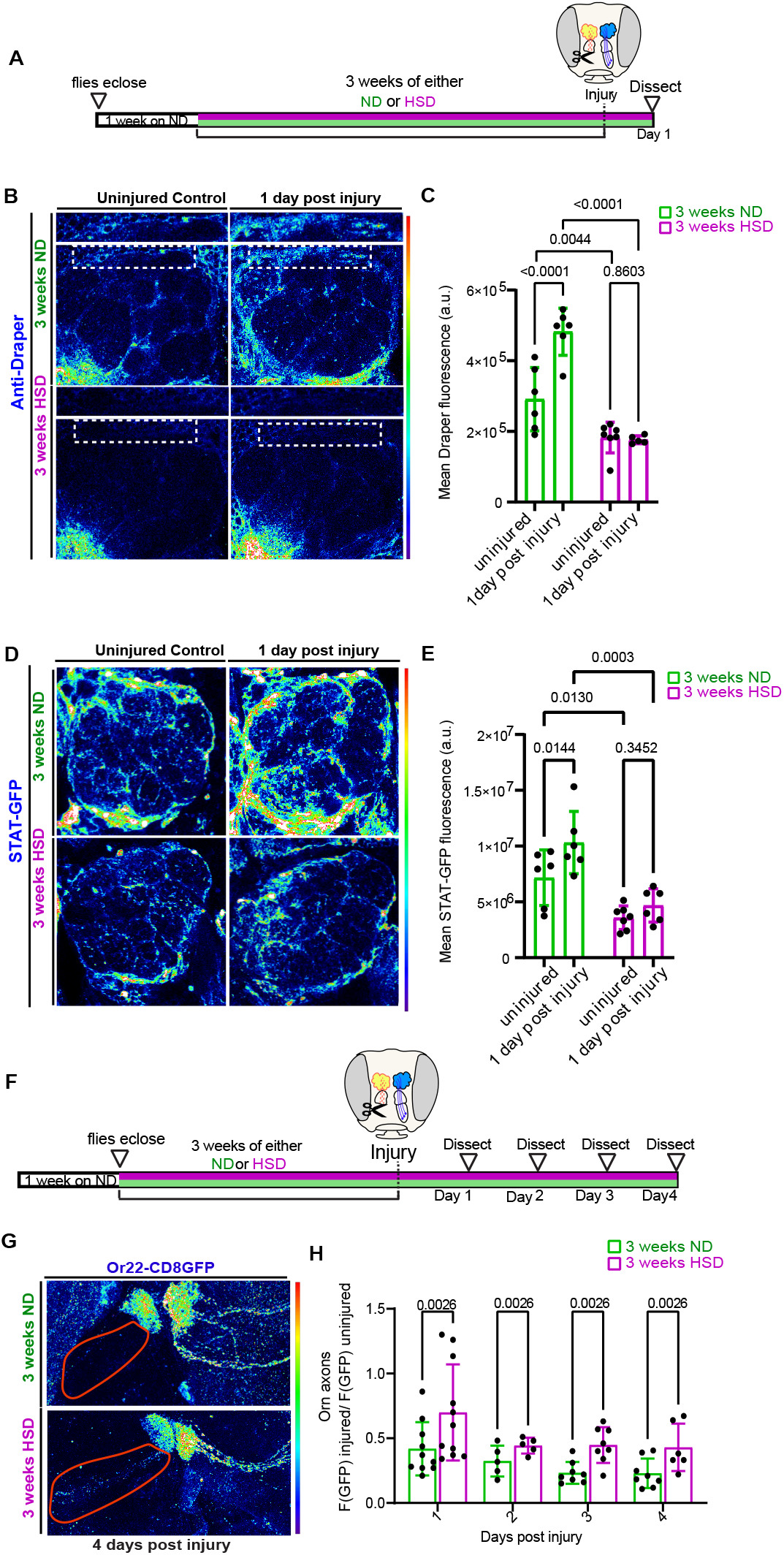
HSD inhibits injury-induced STAT upregulation and delays clearance of degenerating axons. **A)** Schematic showing the experimental strategy for B-E. **B)** Confocal images of Draper levels in the antennal lobe region of uninjured and injured ND or HSD-fed flies. **C)** Mean fluorescent intensity of Draper measured within a region of interest (white box) that coincides with the location of ensheathing glia in ND and HSD-fed flies. **D)** Confocal images of Stat92E levels in the antennal lobe region of uninjured and injured ND or HSD-fed flies. **E)** Mean fluorescent intensity of GFP measured within a region of interest (white box) that covers the antennal lobe in ND and HSD-fed flies. Two-Way ANOVA. N= each circle represents an individual fly. **F)** Schematic showing the experimental strategy for B-C. **G)** Confocal images of the antennal lobe region of flies expressing membrane-tagged GFP in a subset of olfactory neurons. **H)** Mean fluorescent intensity of GFP measured overtime within a region of interest (axons; red outline) that coincides with the location of olfactory axons in ND and HSD-fed flies.

While baseline levels of Draper are regulated by PI3K [19], it has been shown that Stat92E, a transcription factor that acts downstream of both Draper and Insulin signaling is essential for the injury-induced Draper upregulation [19, 24]. Draper-dependent activation of Stat92E creates a positive autoregulatory loop in which Stat92E upregulates the transcription of the *Draper* gene [19]. Given that HSD treatment causes a diminished injury-induced Draper response (Figure 4B-C), we reasoned that Stat92E signaling would be attenuated in the antennal lobe region of the HSD-fed flies. To address this, we used a transgenic reporter line with ten Stat92E binding sites that drive the expression of a destabilized GFP [19]. We found that while ND-fed flies exhibited the expected post-injury Stat92E upregulation, HSD treatment caused a reduction in Stat92E levels at baseline and inhibited post injury upregulation (Figure 4D-E). Together, these data indicate that chronic HSD attenuate the Stat92E/Draper signaling pathway leading to impaired glial phagocytotic function.

To understand how disrupted Draper signaling in the HSD-fed flies affect glial phagocytic function, we subjected flies that express membrane tagged GFP in a subset of olfactory neurons (Odorant receptor 22a) to either 3 weeks of ND or HSD, then performed unilateral antennal ablation and examined the rate of GFP clearance over time (Figure 4F). By normalizing GFP fluorescence on the injured side to the uninjured side of the same animal, we were able to establish an endogenous control (Figure 4G). We found that HSD-fed flies had higher levels of GFP florescence at every timepoint indicating a delay in axonal clearance (Figure 4G-H). Together this data indicate that diet-induced glial insulin resistance impairs the clearance of neuronal debris by downregulating Draper.

## Discussion

With increased life expectancy, age-related neurodegenerative disorders are expected to rise, placing a tremendous burden on the healthcare system. Large-scale epidemiological studies have found that mid-life obesity is an independent risk factor for developing neurodegenerative disorders [1–3]. However, the mechanism underlying this connection remains unclear. Here, using a *Drosophila in vivo* model, we draw a causal link between diet-induced obesity and impaired glial phagocytic function, a major contributor to the pathology of age-related neurodegenerative disorders [65]. We show that excessive systemic insulin signaling leads to glial insulin resistance (Figure 2), which dampens the expression of the engulfment receptor, Draper (Figure 3), resulting in impaired glial clearance of degenerating axons (Figure 4). Together, our study provides a strong mechanistic insight into how diet-induced obesity alters glial function, thereby increasing the risk of neurodegenerative disorders.

### HSD causes glial insulin resistance

Insulin signaling is critical for cell metabolism and function. However, excess systemic insulin can lead to insulin resistance, which results in diminished cellular response. Obesogenic diets are well known to cause insulin resistance in peripheral tissues, including fat, which rely on insulin to regulate glucose uptake[58, 66]. However, the evidence for diet-induced insulin resistance in the brain is scarce. This is mainly due to the belief that the brain is insulin independent [6], though evidence suggests that it acts on neurons and glia in a glucose-independent manner [67].

Our previous work has demonstrated that *Drosophila* can serve as an excellent diet-induced obesity model as its adipose tissue also undergoes insulin resistance when subjected to prolonged HSD treatment [25]. Thus, we used the same dietary regimen to investigate whether obesogenic diets lead to brain insulin resistance. Microglia play a significant role in maintaining nervous system homeostasis and their dysfunction is implicated in a myriad of metabolic and neurodegenerative disorders [62, 68, 69]. Although microglia express the insulin receptor [15], it remains unknown whether obesogenic diets can result in glial insulin resistance. In this study, we showed that prolonged HSD treatment attenuated glial PI3K signaling, a downstream arm of the insulin receptor signaling pathway (Figure 2D-F). This coincided with reduced Ldh levels indicating depressed glycolysis, which is tightly regulated by insulin signaling [42, 44, 45] (Figure 1B-C). Consistent with this, insulin resistance is associated with reduced glycolysis [70, 71]. Interestingly, a prerequisite to microglial activation is the metabolic switch from oxidative phosphorylation to glycolysis [72, 73] Intriguingly, even though glycolysis appears to be disrupted in the brains of HSD-fed flies, we did not observe any changes to glial mitochondrial morphology (Figure 1D-F), a biological response we see in the adipose tissue (Figure 1G-K). This suggest that HSD-induced insulin resistance may prevent glial activation by disrupting their OxPhos-Glycolysis balance. Because Ldh expression and mitochondrial morphology are indirect readouts of glycolysis and OxPhos, respectively, more precise metabolic analysis is needed to determine the specific glial metabolic defects caused by HSD. Furthermore, it is important to acknowledge that both the PI3K and Ldh reporters are non-cell specific. Therefore, it is likely that HSD is causing a downregulation of PI3K signaling and glycolysis in both glia and neurons. Given that glia and neurons are intricately connected, it is possible that dysfunctional neuronal metabolism further exacerbates glial dysfunction.

There are seven *Drosophila* insulin like peptides (Dilps). Dilps 2, 3, and 5 are primarily produced by the IPCs, which reside in the pars intercerebralis (PI) region of the *Drosophila* brain-the invertebrate equivalent to the mammalian hypothalamus. In the adult fly brain, the IPCs terminate their axons on peripheral targets including the gut and aorta for systemic Dilp release[49, 74, 75]. It is possible that excess circulating Dilps induced by HSD cause glial insulin resistance indirectly by dysregulating peripheral organs, such as muscles, guts, and adipose tissue. Nevertheless, distinct IPC arborizations have been observed in the brain, specifically in the tritocerebrum proximal to the antennal lobes, raising the possibility that the IPCs act in a paracrine manner. Given that genetic activation and inhibition of the IPCs for a short period of time was sufficient to influence glial PI3K signaling (Figure 2), it is possible that IPC-released Dilps act directly on ensheathing glia. In the future, it would be interesting to untangle the IPC-ensheathing glia insulin signaling circuit.

### HSD-induced glial dysfunction resembles that caused by aging

A hallmark of neurodegenerative disorders is the failure to clear neuronal debris and cytotoxic proteins, triggering a cascade of devastating effects that include inflammation, cell death, and impaired regeneration. Therefore, it is not surprising that microglial dysfunction is implicated in driving the pathogenesis of many neurodegenerative disorders [12]. Although it is known that microglia express the insulin receptors and respond to insulin treatment *in vitro* [16, 76], it remained unknown whether they experience insulin resistance and whether that impacts their phagocytic activity. Both obesity and age-related neurodegenerative disorders are associated with dysfunctional insulin signaling [58, 66, 77, 78], suggesting a potential link. Here, we show that diet-induced insulin resistance disrupts glial phagocytic activity (Figure 4G-H) by downregulating the phagocytic receptor, Draper (Figures 3 & 4). We were able to demonstrate the direct effects of insulin signaling by showing that Dilp release alone mimicked HSD-induced glial defects (Figures 2 & 3D-L). While other groups have shown that local glial insulin receptor activity regulates Draper expression [24], we found that systemic insulin signaling directly regulates glial Draper expression.

It has been demonstrated by other groups as well as us that physiological factors affect glial function in *Drosophila*. Stanhope and colleagues[79] found that sleep plays a crucial role in glial phagocytosis. Similarly to HSD treatment, sleep loss causes Draper to downregulate, which leads to a failure to clear neuronal debris after injury. As with obesity and type 2 diabetes, dysregulated sleep has been associated with neurodegenerative disorders, further supporting the link between dysfunctional glial phagocytosis and neurodegenerative disorders. It is interesting to draw parallels between the results of this study and a recent study by Purice and colleagues [20] that found that aged flies exhibit delayed axonal clearance because of impaired Draper and PI3K activity. The connection between HSD exposure and aging holds true for humans and mammals as well. Age and neurodegenerative disorders are associated with reductions in MEGF10 [80], the mammalian ortholog of Draper, impaired glial phagocytosis [13], and stunted glycolysis [81]. One study showed that Insulin infusion into young rats activated microglia, but this effect was not observed in older rats suggesting that microglia’s insulin sensitivity is age dependent [82]. Furthermore, similar to our findings, which show an accumulation of lipids in the brains of HSD-fed flies, aged microglia also accumulate lipid droplets leading to impaired phagocytic function [41]. As a result, it could be argued that chronic HSD exposure may “accelerate” aging in flies. It would be interesting to explore this connection in more detail by comparing the transcriptomes of aged and HSD-treated flies in the future.

## Methods

### Fly husbandry

The following *Drosophila* strains were used: Or22a-mCD8GFP (BDSC #52620), Ensheathing glia-Gal4 (BDSC # 39157), 10xSTAT92E-GFP (Perrimon Lab), LDH-GFP (a generous gift from Dr. Tennessen), tGPH (Gift of Bruce Edgar), Dilp2-Gal4 (a gift from P.Shen), UAS-TrpA1 (BDSC #26263), UAS-EKO22 (BDSC #40974), UAS-PI3K-CAAX (BDSC #8294). Flies were housed in 25°C incubators and all experiments were done on adult male flies. To induce the expression of the temperature sensitive TrpA1, flies were moved to a 29°C incubator 1 week post eclosion. Following 1 week after eclosion, the flies were placed on either a normal diet containing 15 g yeast, 8.6 g soy flour, 63 g corn flour, 5 g agar, 5 g malt, 74 mL corn syrup per liter, or a HSD, which consists of an additional 300 g of sucrose per liter (30% increase).

### Antennal nerve injury

As adapted from [19–21, 23, 24, 64, 79, 83, 84], flies were anesthetized using CO2 and antennal nerve injury was accomplished by unilaterally removing the third antennal segment of anesthetized adult flies using forceps. Flies were then placed back into either ND or HSD until they were dissected 24 hours after injury or as indicated otherwise in the figure legends.

### Immunostaining

Immunostaining of adult brains and fat bodies was performed as previously described[25, 50, 51]. Tissues were dissected in ice-cold PBS. Brains were fixed overnight in 0.8% paraformaldehyde (PFA) in PBS at 4°C. The fixed brains were washed five times in PBS with 0.5% BSA and 0.5% Triton X-100 (PAT), blocked for 1 hour in PAT + 5% NDS, and then incubated overnight at 4°C with the primary antibodies. Following incubation, the brains were washed five times in PAT, reblocked for 30 min, then incubated in secondary antibody in block for 4 hr at room temperature. Finally, the brains were washed five times in PAT, then mounted on slides in Slow fade gold antifade. Primary antibodies were as follows: rabbit anti-Dilp5 (1:500; this study); Chicken anti-GFP (1:500; Cat# ab13970, RRID:AB_300798); And Mouse anti-Draper (1:50; DSHB 5D14 RRID:AB_2618105). Secondary antibodies from Jackson ImmunoResearch (1:500) include donkey anti-Chicken Alexa 488 (Cat# 703-545-155, RRID: AB_2340375); donkey anti-mouse Alexa 594 (Cat# 715-585-150, RRID:AB_2340854). Lipid droplets were stained with lipidtox (1:500, Thermo Fisher Cat#H34477) overnight at 4C.

### Image analysis

Images were acquired with a Zeiss LSM 800 confocal system and analyzed using ImageJ[85]. All images within each experiment were acquired with the same confocal settings. Z-stack summation projections that spanned the depth of the antennal lobes at 0.3 um intervals were generated and a region of interest (indicated on the figure) was used to measure the fluorescent intensity of GFP or Draper. Dilp 5 was measured using z-stack summation projections that included the full depth of the IPCs. A region of interest around the IPCs was manually drawn using the free hand tool and the integrated density values were acquired. To measure mitochondrial morphology, A maximum-intensity projection of Z-stacks that covered the full depth of the antennal lobe was used for ImageJ analysis. The maximum-intensity projection was first inverted then automatically thresholded before the ‘analyze particles’ function was used to measure the average mitochondrial circularity, major axis, and minor axis.

### statistics

All the statistical tests were done using GraphPad PRISM. (GraphPad Software Incorporated, La Jolla, Ca, USA, RRID:SCR_002798). The assumption of normality was tested using Shapiro-Wilk’s test. A *t* test was performed for Figures 1–3, while a 2-way ANOVA was used for Figures 4–5. Data are presented as means ± standard deviation. Significance was set at p<0.05. N for each experiment is detailed in the figures.

## Acknowledgments

We thank Dr. Jason M Tennessen for generously donating the Ldh-GFP transgenic fly line used in this article. This work is possible due to grants awarded to AR from the NIGMS (R35GM124593) and the Brain Research foundation (BRFSG-2022-09). Mroj Alassaf is supported by a postdoctoral fellowship from the Helen Hay Whitney Foundation.

---

**Figure S1.**
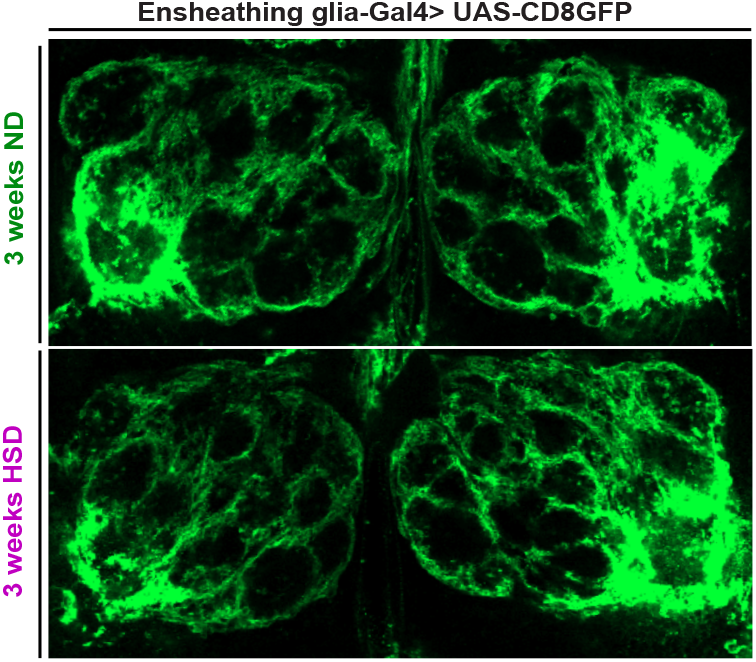
HSD does not affect glial gross morphology. The antennal lobe regions of flies with ensheathing glia-driven membrane tagged GFP. No observable gross morphological defects in the HSD-fed flies.

